# VisEgg: A robust phenotyping tool to assess rainbow trout egg features and viability

**DOI:** 10.1101/2020.04.21.052472

**Authors:** Emilie Cardona, Jérôme Bugeon, Emilien Segret, Julien Bobe

**Author notes:** equal contribution (Emilie Cardona, Jérôme Bugeon).

## Abstract

Assessing female fish reproductive success requires a thorough evaluation of egg characteristics, including egg number, size and variability as well as egg developmental potential through the monitoring of embryo survival after fertilization. While embryonic success relies, at least in part, on paternal contribution, some parameters are strictly related to egg characteristics, one of the main ones being the viability of the egg when released into the water at spawning. It is however not necessarily possible, at least in salmonid fish that lay non-transparent eggs, to separate the different causes of egg/embryo failure.

In this context, our aim was i) to develop a simple and rapid system to capture images of rainbow trout eggs combined with computerized processing of these images to perform a fully automatic individual characterization of egg features including number and size ii) to estimate unfertilized egg viability through the monitoring of the percentage of eggs that will not survive to water hydration.

To evaluate the VisEgg system, unfertilized eggs (approximatively 400 eggs per batch) originating from 105 different females were hydrated in water. After 24h, a picture of the eggs was obtained using a dedicated shooting system consisting of a light source and a digital single-lens reflex (SLR) camera. An image processing algorithm was developed to allow the automatic detection and separation of the eggs and to perform automatic measurements of egg number and individual egg size. The presence of white egg was used as an indirect measure of egg integrity, the “whitening” being the result of water entry into the egg through the vitelline membrane. These white eggs were therefore considered as non-viable, as a result of their lack of physical integrity.

Fertilization assays were performed in parallel using a subsample of the same egg batch. Embryonic development was monitored and hatching rate was calculated. A significant correlation between white egg percentage after hydration and hatching rate was observed (Spearman coefficient =-0.557, p<0.001), in consistency with the fact that non-viable egg will not allow successful embryonic development. In contrast, the percentage of eggs that do not successfully hatched includes egg/embryo failures of different nature including egg viability, their capacity to be fertilized and to develop into an embryo. Using VisEgg, we were able to quantify the lack of viability of the eggs separately from the different other events that may occur during fertilization and incubation. VisEgg is a convenient and reliable tool to obtain individuals measures on trout eggs. It can be used to assess not only egg size and egg number but also unfertilized egg viability before fertilization.

## Introduction

The control of egg quality (i.e. the ability of the egg to be fertilized and subsequently develop into a normal embryo) is of major importance for many, if not all, aquaculture fish species (Migaud et al., 2013). A regular supply of high quality eggs is mandatory for rainbow trout hatcheries sustainability (Bromage et al., 1992). Given the high cost of broodstock rearing, large variations in the quantity or quality of gametes can significantly impact the competitiveness and sustainability of fish farms and aquaculture companies (Bromage et al., 1992, Bobe, 2015). Hence, controlling egg quality is a major issue in aquaculture with important economic consequences.

Teleost eggs consist of an inner ooplasm comprising vitellus (or yolk) surrounded by the vitelline membrane also known as plasma membrane. The outer membrane called the chorion is a thick layer with a single micropyle. Spermatozoa must pass across the micropyle to penetrate the egg and allow fertilization (Kuchnow and Scott, 1977). Teleost eggs are soft in the ovary and when the egg is freshly stripped from the female, the chorion is limp and the egg is flaccid, egg is not really round. A thin space between the chorion and the vitelline membrane exists and corresponds to the perivitelline space. When eggs are transferred into water, a contractile activity of the yolk mass occurs as a result of subsequent discharge of the cortical vesicle contents in the perivitelline space (Kobayashy, 1985). Colloid substances, with water taking up property, are thus discharged into the perivitelline space. Water is osmotically drawn through the fine pores of the chorion into the perivitelline space. This process creates a turgid pressure in the egg and the chorion is stretched until a steady state between internal and external environment is achieved, resulting in the hardening of the chorion (Alderdice, 1988). The stage of equilibrium is reached in about 20 minutes. This phenomenon is called egg water hardening process and egg from rainbow trout absorbs approximatively 20% of its initial volume in water (Yamamoto 1962; Blaxter 1969; Lahnsteiner et al. 1999). The vitelline membrane is the only significant barrier to the diffusion of water and ions between the vitellus and the external medium. When eggs are of good quality, the vitelline membrane is highly impermeable to water and electrolytes and it is not easy to cause a rupture of the membrane (Gray, 1932). Inversely, when eggs are of bad quality, the vitelline membrane is weakened and the water pressure is sufficient to break it. The vitellus is rich in vitellogenin and protein, specifically in globulin. Globulins and vitellogenins are soluble due to the salts contained in the vitellus. But when the membrane breaks, the water enters and salts are diluted resulting in the precipitation of globulins (Gray, 1932; Van Heerden et al., 1996) and vitellogenins (Engelmann et al., 1976; Fremont and Riazi, 1988). In this case, egg turns white. The presence of white egg was thus interpreted as an indirect measure of vitelline membrane integrity, the “whitening” reflecting the result of water entry into the ooplasm. These white eggs were thus considered as non-viable.

Improvement of female reproductive traits (egg diameter, absolute fecundity, number of viable eggs) through changes in their rearing conditions (feed, temperature, light cycle, etc.) or genetic selection implies an ability to measure the quality of the spawn in a very large number of individuals. The assessment of fish breeding performance requires, in particular, a precise evaluation of the number of spawned eggs but also information on the egg ability to be fertilized and subsequently develop into a viable embryo. Prior to fertilization, it is extremely difficult to estimate differences between good and bad quality eggs in rainbow trout, despite the fact that the characterization of predictive estimators or egg quality markers would have major applications for research and industry (Migaud et al., 2013). For the above reasons, enabling a comprehensive evaluation of egg quality before fertilization is highly desirable in aquaculture. Moreover, during fertilization and incubation, poor egg quality can lead to several problems including insufficient egg hydration, lack of fertilization, developmental arrests, embryonic mortalities and deformities (Brooks et al., 1997; Bobe and Labbé, 2010; Migaud et al., 2013). Given the wide variety of the problems observed, it is usually very difficult to determine the causes of embryonic/developmental failure.

In the present study, an automatic phenotyping system based on image analysis was developed to characterize different egg features (number, size) and the occurrence of non-viable eggs in order to separate this cause from other events that may occur later during fertilization or embryonic development. The phenotype measured here is based on the egg ability, specifically the vitelline membrane capacity, to resist to water pressure during hydration process. This method is complementary, yet much more rapid, of a full evaluation of developmental success that is performed on fertilized eggs to separate the origin of egg quality defects (viable eggs vs. non-viable eggs that do not allow developmental success).

## Material & methods

### Ethical statements

Experimentations were conducted in the INRA PEIMA experimental facility (Sizun, France - Agreement number B29-277-02). All fish were reared and handled in strict accordance with French and European policies and guidelines of the INRA PEIMA Institutional Animal Care and Use Ethical Committee, which specifically approved this study. Fish were monitored daily during the experiment. If any clinical symptoms (i.e. morphological abnormality, restlessness or uncoordinated movements) were observed, fish were sedated by immersion in MS-222 solution at a concentration of 50mg.L^−1^ and then euthanized by immersion in a MS-222 solution at a concentration of 400mg.L^−1^ (anesthetic overdose) during 3 minutes.

### Broodstock breeding and experimental design

Female rainbow trout from an autumn-spawning strain were held under natural photoperiod until their first reproduction (2 years) in INRA PEIMA experimental facility. After spawning, fish were transferred into 6 outdoor 2m^3^ tanks (initial weight: 1693 ± 306g). Females were fed a commercial trout broodstock diet (Le Gouessant, Lamballe, France) for a period of five months. During this five-month period, an artificial photoperiod regime was applied to obtain a second reproduction during summer: a 2.5-month long photoperiod (20 hours light, 4 hours dark from December to March, 2017) followed by a 2.5-month short photoperiod (8 hours light, 16h hours dark from March to June, 2017). Water temperature during experiment ranged from 6 to 12°C.

### Sample acquisition

During spawning season (June 21^th^ to August 3^rd^, 2017), females were checked for ovulation once a week by applying a manual pressure onto the abdomen. When ovulation was detected, fish were given an intraperitoneal injection of an antibiotic (vetrimoxin) to prevent *Flavobacterium* infection. Two days after the detection of ovulation, mature females were manually stripped. Eggs were weighted and two samples were taken for fertilization assays and for VisEgg phenotyping analysis. A total of 105 spawns were analyzed individually.

### Fertilization assays

For each spawn (i.e. each female, n=105), approximately 400 eggs were fertilized with a pool of sperm collected from males fed *ad libitum* on a commercial diet and embryonic success was monitored. Fifteen microliters of a pool of semen obtained from five males presenting the highest sperm motility were used, and 15 ml of Actifish solution (sperm motility activating saline solution, IMV technologies, L’Aigle, France) were added onto the eggs. Five minutes later, the sperm motility activating solution was drained and egg batches were transferred into individual incubators in a recirculated water unit. Water temperature (12.0-12.6°C) was monitored daily. Dead eggs and embryos were periodically manually counted and removed. Survival at eyeing, hatching and completion of yolk-sac resorption (YSR) were monitored and were calculated as a percentage of the initial number of eggs used for fertilization. The occurrence of noticeable morphological malformations (spinal cord torsion, head or caudal fin malformations, etc.) at YSR was also recorded.

### Phenotyping analysis

For VisEgg phenotyping, unfertilized eggs (20 to 25 grams corresponding to approximatively 400 eggs) were stored in 100 ml container and then water from the incubation system was added to hydrate the eggs (80mL). After a 24-hour hydration at 4°C, a picture of the egg sample was taken using a dedicated commercially available shooting system. This system consisted in a light tablet and a digital SLR camera (canon EOS 1000D, resolution: 10.1 M pixels) (Figure 1). Eggs were rinsed with water and delicately poured into a petri dish (diameter=14cm). The image calibration was performed with an image of 1 euro coin with a known area to calculate the pixel size. The petri dish was then placed on the light tablet. It was verified that the eggs were not superimposed and a picture was taken. An image processing algorithm was developed and coded using a Visual Basic for Applications (VBA) macro with Visilog 7.3 software (Thermo Scientific). This software allows the automatic detection and separation of the eggs in order to make fully automatic measurements of egg number and specific measurement for each egg (size, shape, color, white egg occurrence). The total number of ovulated eggs per females could be back calculated using the number of measured eggs and the total weight of the spawn. The workflow of the VisEgg macro-command is presented in Figure 2. All details and relevant information about the VisEgg image analysis workflow are available here: https://github.com/LpgpImage/VisEgg/wiki.

**Figure 1:**
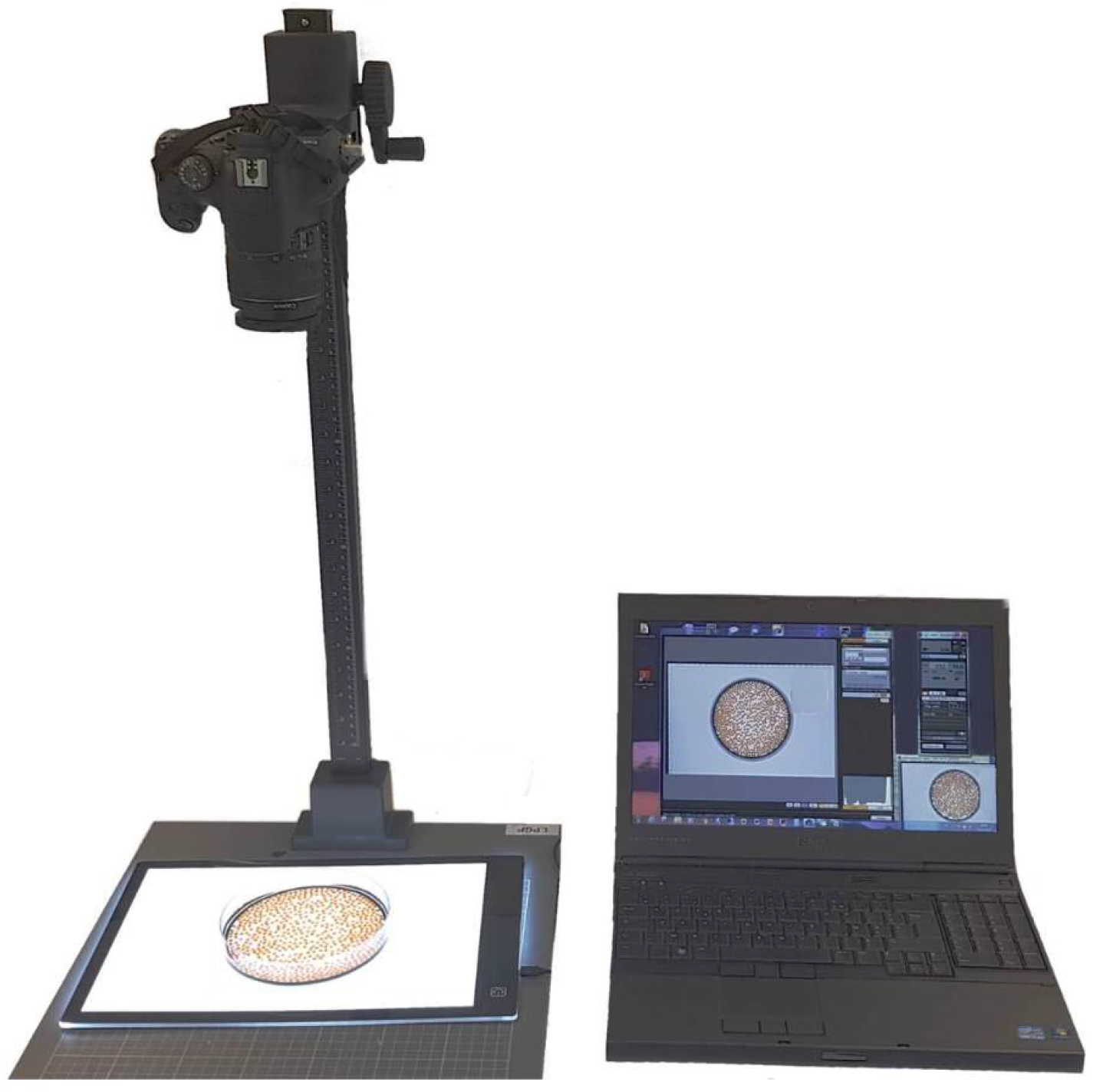
The VisEgg system equipment and setup.

**Figure 2:**
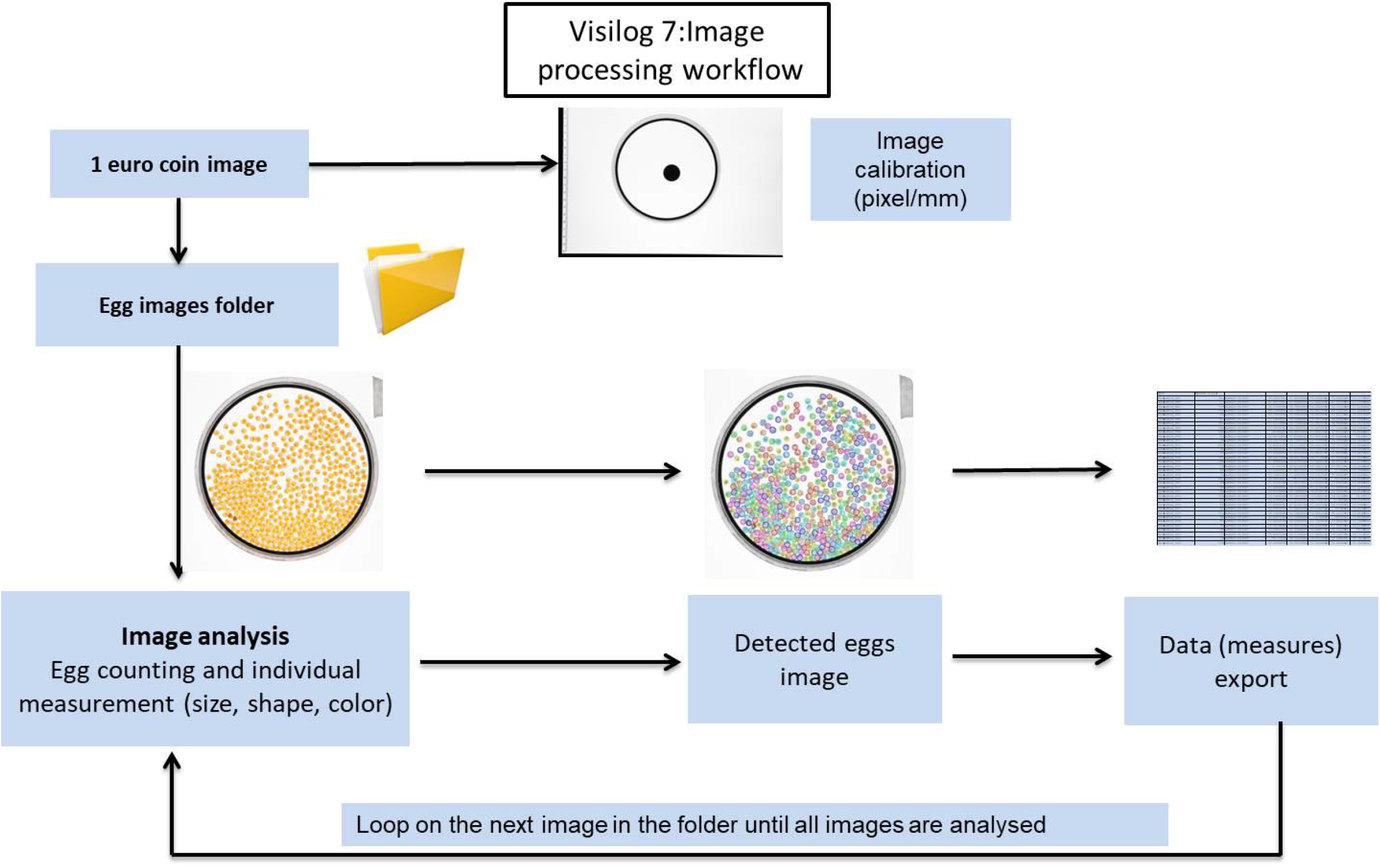
The VisEgg workflow.

### Statistical analysis

Statistical analysis of the data was carried out using the R software. Percentage data were transformed using an arcsine transformation before analysis. Simple linear regressions were performed to test the correlation between survival rates at different stages. Multiple linear regressions (MLR) were performed to test the correlation between different egg quality parameters measured with VisEgg (i.e. egg diameter, variability of egg diameter and white egg percentage) and hatching rate. MLR was simplified to the minimal adequate model by stepwise removing of nonsignificant elements. The adequacy of the final linear regression has been assessed by making residual plots to check the normality. This showed that nothing was amiss; the analysis was thus validated. Spearman coefficients were calculated.

For statistical analysis performed to compare the two calculating methods of hatching rate (see Figure 5), the normality of data distribution and homogeneity of variance were checked. These conditions were not respected; a Kruskall-Wallis test was used followed by Dunn test’s post hoc analysis.

## Results & discussion

### Automatic assessment of egg features

Evaluating the reproductive performance traits requires counting the eggs and monitoring developmental success. This time-consuming task, performed manually in hatcheries, is particularly tedious. In most cases eggs are weighed and not counted, as the weight is easier to determine. Furthermore, measuring the mean weight of a spawn does not provide information on individual variability within each spawn, which is also an indicator of spawn quality. Using the VisEgg automatic phenotyping tool presented here, we were able to observe differences that would have been overlooked using more simple approaches.

The total egg number is indicator of reproductive performance variability between females as illustrated in Figure 3 for two different females: total egg number can be similar despite a different weight of the spawn. When only the total weight of the spawn is used, the size of the eggs remains unknown. Using VisEgg, it is now possible to measure a large number of eggs very easily. In this study, a very large number of eggs (n=52,000 eggs originating from 105 spawns) were automatically individually measured. This significantly increased the accuracy of the measurement. Using VisEgg we were able to measure individual egg size of all sampled eggs and therefore to estimate intra-spawn variability (i.e. individual egg variability) and inter-spawn variability (i.e. individual female variability).

**Figure 3:**
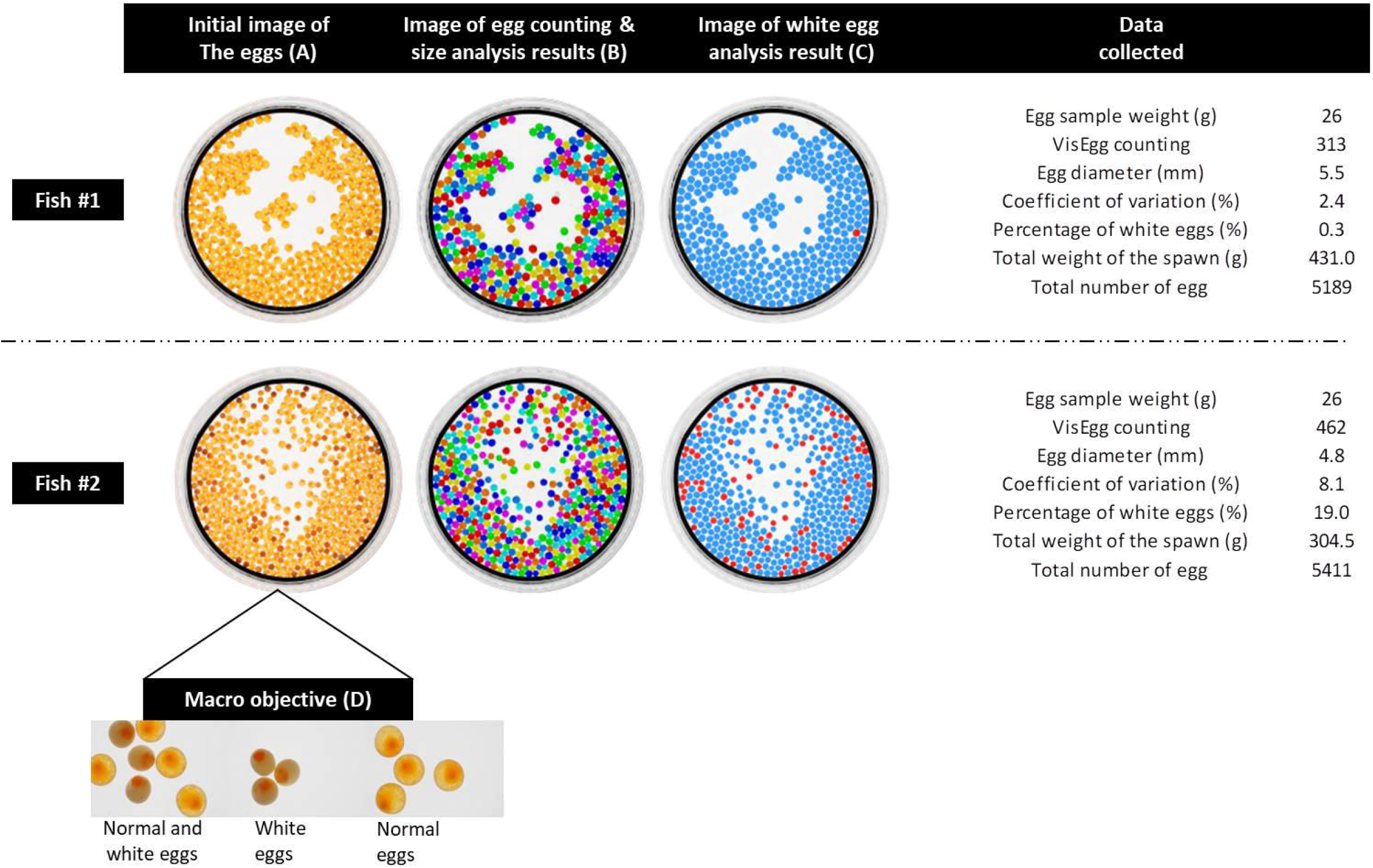
Images of trout eggs as processed by VisEgg. (A) Initial image obtained with VisEgg system equipment presented in Figure 1; (B) Image as processed by VisEgg for an automatic detection and separation of the eggs in order to make fully automatic measurements of egg number and size; (C) Image as processed by VisEgg for an automatic detection of white eggs and normal eggs (white eggs and normal eggs are represented in red and blue color, respectively); (D) Pictures with macro objective illustrating normal and white eggs after hydration.

The distribution of egg diameter obtained with 105 spawns individually analyzed showed values ranging from 3.8 to 5.2 mm with a mean of 4.6 mm. The distribution of the coefficient of variation of egg diameter for all spawns ranged from 2.2 to 9.2% with a mean of 4.6%. These values show a high variability between the spawns analyzed in this study that could not have been observed with a standard technique.

Egg size remains an interesting parameter to measure. Although the possible benefit effect of egg size on egg survival and alevins is still debated, especially under aquaculture conditions (Bobe and Labbé, 2010; Bromage et al., 1992; Campbell et al., 1992; Jastrebski and Morbey, 2009; Migaud et al., 2013; Springate and Bromage, 1985b), it is established that larger eggs produced larger alevins (Springate and Bromage, 1985) with substantial fitness advantages over small alevins (Heath et al., 2003). While egg size is very often measured in studies on fish reproductive performance, it is unusual to observe egg size variability in intra spawn as a measured parameter. This is certainly due to the difficulty of assessing this parameter. Female rainbow trout (*Oncorhynchus mykiss*) is a group-synchronous species which produces a single spawn each year where all oocytes develop and ovulate at the same time (Lubzens et al., 2010). In normal conditions, eggs from a single spawn are of homogeneous size; the presence of large egg size variability in intra-spawn for salmonid can be the consequences of a disruption of the physiological processes underlying oogenesis (Jastrebski and Morbey, 2009). The measurement of this parameter therefore appears to be relevant for rainbow trout reproduction studies.

Thus, VisEgg is a convenient and reliable tool to obtain accurate individuals measures, as egg size and variability.

### Assessment of egg viability regardless of fertilization and subsequent developmental success

Unfertilized eggs rather than fertilized eggs were used to evaluate our phenotyping tool because hardening process is independent of fertilization (Lahnsteiner et al., 1999). Two distinct phenotypes can be observed after the hydration process. Either the egg hydrates normally and grows slightly or the egg turns white. This phenotype is easily observable on images (Figure 3D). Moreover, VisEgg can separate normal (i.e. viable) eggs from white (i.e. non-viable) eggs and then calculate the percentage of white egg for each analyzed egg batch (Figure 3C).

In this study, multiple linear regression was performed to test the correlation between hatching rate and the different parameters measured with VisEgg (i.e. egg diameter, variability of egg diameter and white egg percentage). Finally, only white egg percentage was significantly correlated with hatching rate. Here, egg size and variability appear to have no link with egg quality and alevins survival (Spearman coefficient=0.39 and −0.30, respectively for egg diameter and variability of egg diameter). Linear regression between white egg percentage and hatching rate shows a significant negative correlation (p-value=2.6 x10^−10^; Spearman coefficient=-0.56).

It should be noted that we observed highly significant correlations between surviving rates at eyeing and hatching as well as between surviving rates at hatching and yolk-sac resorption (*p-value*= <2 x10^−16^, Spearman coefficient=-0.99 for both correlations). Surviving rates at eyeing and YSR are therefore also significantly correlated with the percentage of non-viable eggs (*p-value*=9.9 x10^−11^, Spearman coefficient=-0.58 and *p-value*=4.7 x10^−10^, Spearman coefficient=-0.55 for eyeing and YSR, respectively).

The negative correlation between white egg percentage and developmental success was expected because non-viable eggs will not allow successful embryonic development. In addition, the modest coefficient of correlation can be explained by the composite nature of survival rate that includes the capacity of egg to survive in water (i.e. egg viability), to be fertilized and to develop into an embryo (or to die at different times during development). Figure 4 illustrates the successive different origins of egg/embryo failure between spawning and yolk-sac resorption. The graph shows means values obtained using the 105 different spawns analyzed in the present study.

**Figure 4:**
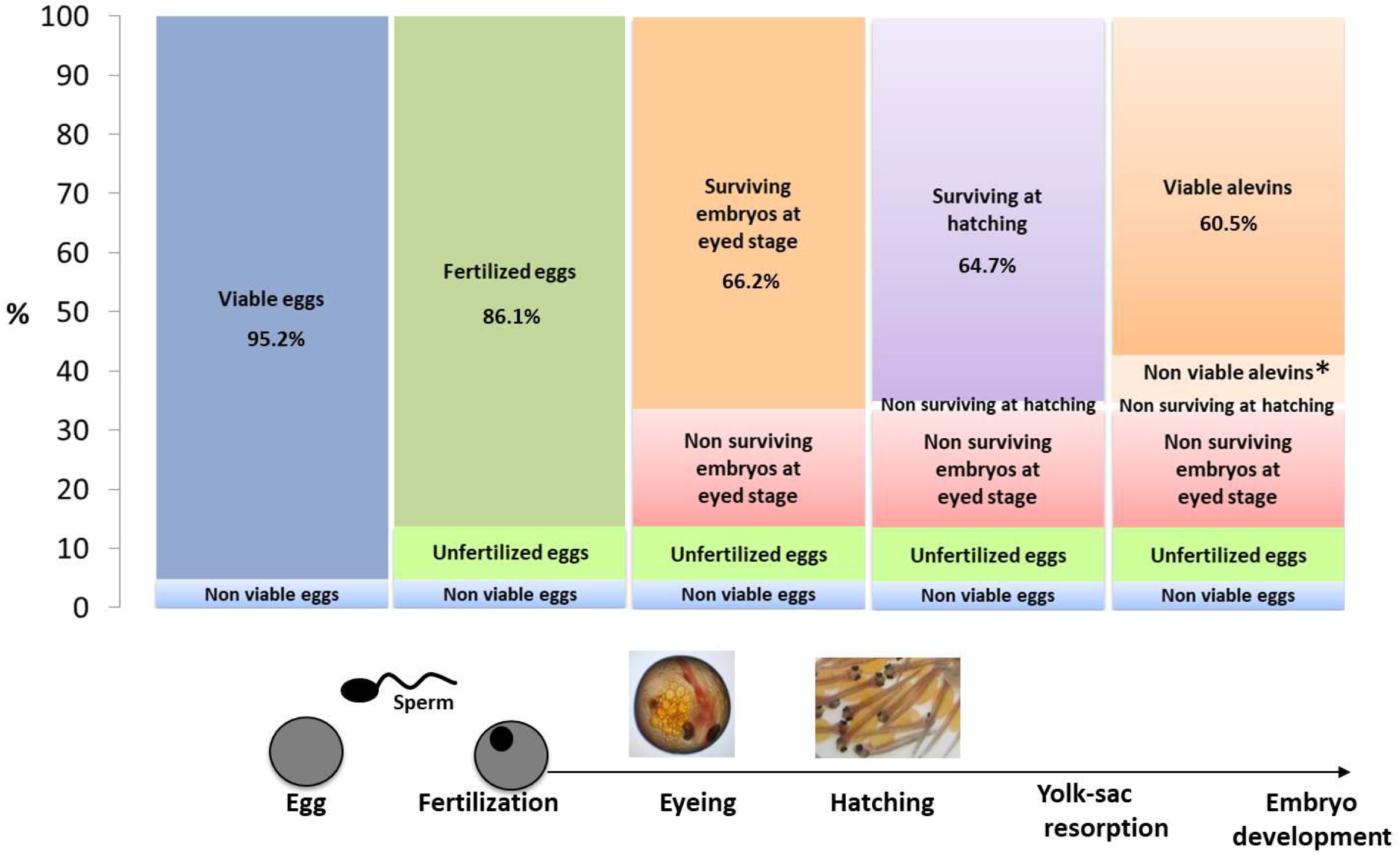
Different causes and relative importance of egg quality defects calculated using VisEgg and embryonic monitoring. Data represent the average percentage values calculated from 105 egg batches followed individually. Viable and non-viable eggs data were obtained with VisEgg system. Fertilized eggs and survival at eyeing, hatching and YSR are expressed as a percentage of the initial number of eggs used for fertilization. Non-malformed alevins with full completion of yolk-sac resorption are considered as viable alevins. Alevins with noticeable morphological malformations (spinal cord torsion, head or caudal fin malformations, etc.) and dead alevins between hatching and resorption are considered as non-viable alevins (*non-viable or malformed alevins).

Figure 5A shows the hatching rate calculated using all eggs (in blue) or only on viable eggs (in red), the batches being ranked based on the overall hatching rate (blue dots). The blue and red dots do not follow exactly the same pattern due to differences in the origins of egg/embryo failure up to hatching, with major differences between the two groups for some individuals. In some cases, when the red and blue dots are far apart, there is a high incidence of egg viability on the overall developmental success. In contrast, when the red and blue dots overlap (or are close) there is a low incidence of egg viability on the overall developmental success. Figure 5 (A&B) shows that a high incidence of egg viability is observed for a significant number of females. It should be noted that the importance of egg viability is highly variable as illustrated by significant differences between red and blue dots that can be observed regardless of the overall development success with the exception of high quality spawns (> 80% hatching rate; Figure 5B). In 15.2% of the cases, there is more than 10% of difference between red and blue dots. The percentage of batches over 10% variation between the two calculating methods of hatching rate can reach 50% of the batches for hatching rates ranging from 21 to 40%. While it is not necessarily easy to decipher the causes of egg/embryo failure up to hatching, VisEgg offers the possibility to quantify the percentage of viable eggs. As just an example, four particular cases are shown in figure 6 to illustrate the different causes that can lead to a similar developmental success. Fish A and Fish B present low and equivalent hatching rates (23.0% and 24.3%, respectively). However, Fish A exhibits a high incidence of egg viability (39.6%) on the overall developmental success whereas Fish B exhibits a low incidence of egg viability (0.2%) on the overall developmental success. Similarly, Fish C and Fish D exhibit high hatching rates (65.2% and 69.2%, respectively) but for different reasons: Fish C exhibits a high incidence of egg viability (22.1%) on the overall developmental success whereas Fish D exhibits a low incidence of egg viability (1.1%). These two examples clearly illustrate that similar developmental success (either low or high) can be observed even though the phenotype, and most likely the biological underlying cause, is completely different. The VisEgg tool offers the possibility to separate and quantify the incidence of non-viable eggs on the overall developmental success rather than loosing this information when only developmental success is monitored. This approach is thus complementary of a full evaluation of developmental success that is performed on fertilized eggs to separate the origin of egg quality defects (viable eggs vs. non-viable eggs that do not allow developmental success).

**Figure 5:**
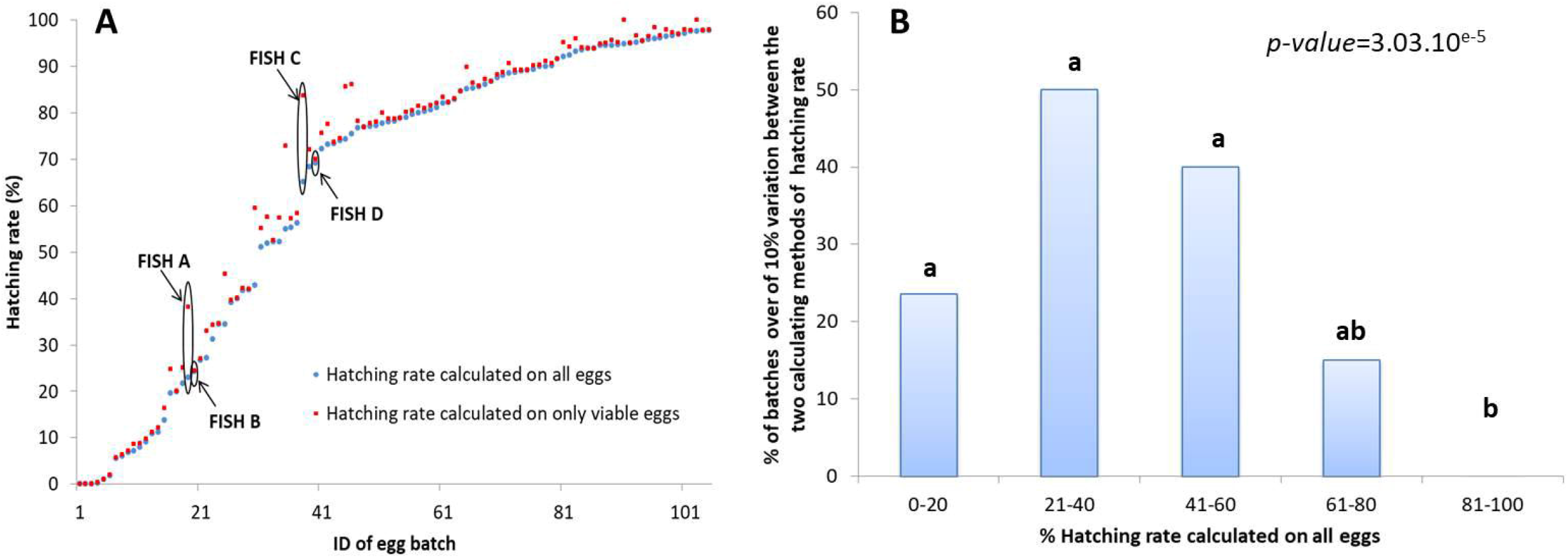
(A) Hatching rate calculated on number of all eggs (blue) or on only viable eggs (red); (B) Percentage of batches with ≥10% variation between the two calculating methods of hatching rate. Statistical differences were evaluated by Kruskal-Wallis test followed by Dunn test post hoc analysis. Different causes of egg quality defects for FISH A, B, C and D were represented in figure 6.

**Figure 6:**
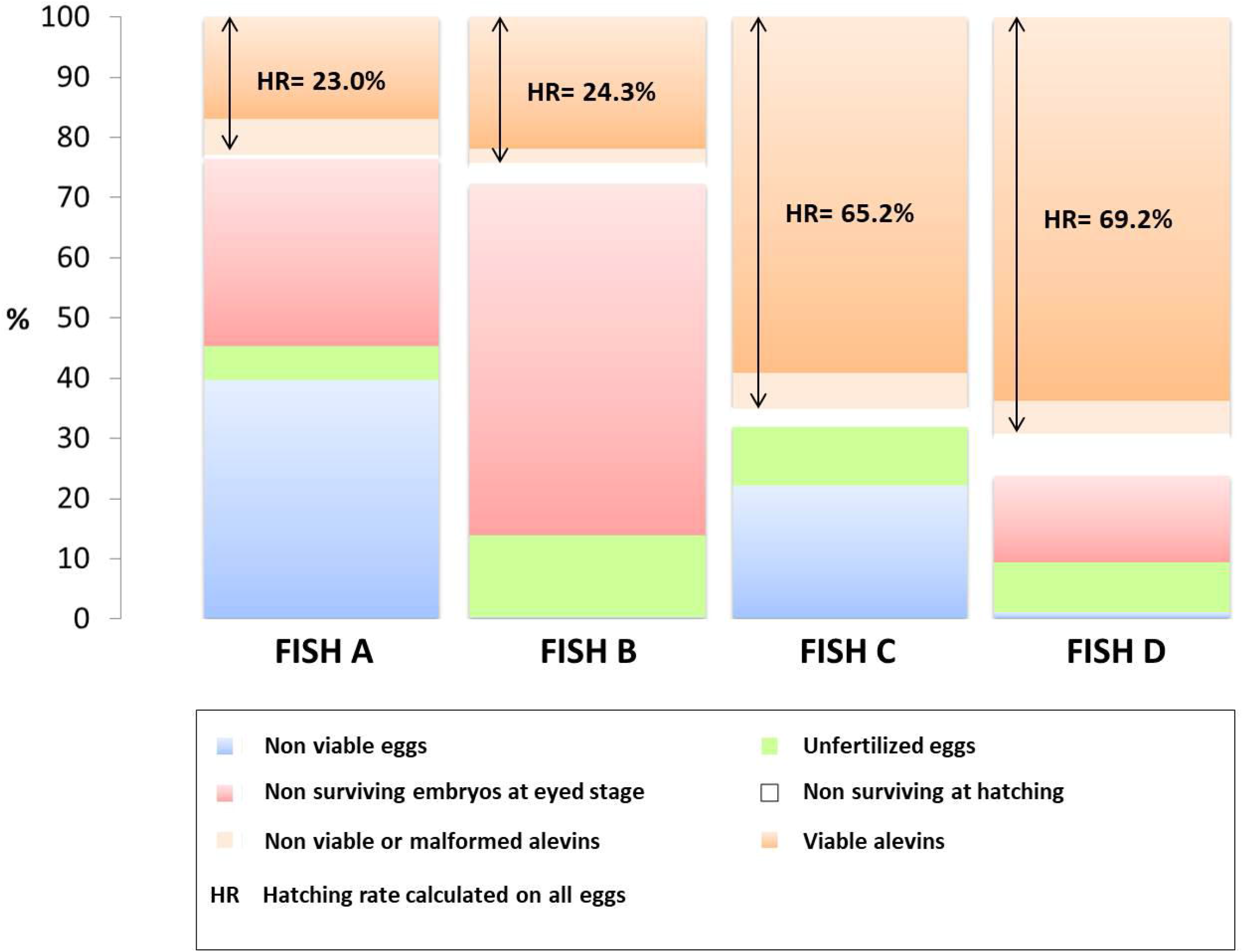
Different causes of egg quality defects that can occur between spawning and yolk-sac resorption in four fish displayed on figure 5. See legend from figure 4 for details on calculation methods.

## Conclusion

In summary, VisEgg is a convenient, fast, automatic and reliable tool to obtain individual measures of trout eggs. It can be used to assess not only egg size and number but also to assess unfertilized egg viability before fertilization. VisEgg is the first automatic phenotyping tool for egg viability, a key component of egg quality. This tool, developed in rainbow trout, is now being used routinely on trout eggs but it could be adapted to the eggs of other species such as sturgeon. In the future, VisEgg will be adapted to measure other egg characteristics such as color, shape (eccentricity value) and over ripening eggs.

## Acknowledges

We thank François Guivarc’h for animal care and all the people of PEIMA and LPGP for their help during reproduction season.

---

Supplementary Table 1: Table presenting raw data for all 105 spawns used for the construction of Figure 4 and 5.

